# Stratification of Behavioral Response to Transcranial Current Stimulation by Resting-State Electrophysiology

**DOI:** 10.1101/2020.01.27.921668

**Authors:** Atalanti A. Mastakouri

## Abstract

Transcranial alternating current stimulation (tACS) enables the non-invasive stimulation of brain areas in desired frequencies, intensities and spatial configurations. These attributes have raised tACS to a widely used tool in cognitive neuroscience and a promising treatment in the field of motor rehabilitation. Nevertheless, considerable heterogeneity of its behavioral effects has been reported across individuals. We present a machine learning pipeline for predicting the behavioral response to 70 Hz contralateral motor cortex-tACS from Electroencephalographic resting-state activity preceding the stimulation. Specifically, we show in a cross-over study design that high-gamma (90–160 Hz) resting-state activity predicts arm-speed response to the stimulation in a concurrent reaching task. Moreover, we show in a prospective stimulation study that the behavioral effect size of stimulation significantly increases after the stratification of subjects with our prediction method. Finally, we discuss a plausible neurophysiological mechanism that links high resting-state gamma power in motor areas to stimulation response. As such, we provide a method that can distinguish responders from non-responders to tACS, prior to the stimulation treatment. This contribution could eventually bring us a step closer towards translating tACS into a safe and effective clinical treatment tool.

## I. Introduction

Non-invasive brain stimulation (NIBS) modulates neural activity, behavior, and brain plasticity through the non-invasive creation of forced electrical current flows inside the brain [1], [2]. There are two main categories of NIBS: Transcranial Magnetic Stimulation (TMS), which uses external magnetic fields to force the creation of electrical potentials in the cortex that depolarize neurons and trigger action potentials [3], and Transcranial Electrical Stimulation (TES) [4], which applies weak electrical direct (tDCS) or alternating (tACS) currents on the scalp [5]. In contrast to TMS, only a fraction of this current enters the brain and causes a membrane potential change of the affected neurons. It has been shown by [6] that this membrane potential could be sufficiently strong to change their probability of generating action potentials. Nevertheless, as the physiological effects of tACS are not completely uncovered yet, there is still a debate on this matter, particularly with low intensity currents. Recent publications [7], [8] demonstrated entrainment of neural activity to tACS on primates. For humans, there are some limited evidence for phase-specific entrainment in EEG after-effects for alpha-frequencies. [9], [10].

Although the neural mechanisms of NIBS are not yet fully understood [11], NIBS applications are becoming more widespread in research and treatment. Applications of NIBS can be divided into three main categories: studies that probe neurophysiology (e.g., how neural oscillations are causally related) [12]–[18], studies that investigate how brain activity gives rise to behavior and cognition [11], [19]–[28], and studies that employ NIBS for rehabilitation [29]–[33].

In all three categories, NIBS studies report substantial variations in stimulation response across individual subjects, including up to 55% of non-responders [34], [35]. While non-responders decrease the statistical power of NIBS studies, this sub-group is unproblematic from an ethical point of view. Given the reported variances in effect sizes, however, it is not unreasonable to assume that there also exist subjects with negative stimulation responses [34]–[38]. Here we use the term “negative responders” to identify those subjects that exhibited a response oposite to what was expected which may be harful or not. We do not necessarily imply that this term refers to harmful effects only, although it may include them. Subjects with negative stimulation responses would be highly problematic, because their existence would imply that NIBS studies may violate the principle of not having detrimental effects. This ethical concern is particularly relevant in clinical settings, where NIBS is used to cause long-term changes [39], [40]. In a related work, Yang *et al.* [41] and Kasten *et al.* [42] stress the necessity of pre-stimulation screening and the importance of individualizing stimulation protocols. Furthermore, Kong *et al.* [43] report the existence of individual-specific cortical networks and their importance in the prediction of human cognition, giving evidence for individual cortical differences. Taking all this into account, ethical use of NIBS would first require consideration of potential negative stimulation effects in individual subjects, and, second, if such effects exist, implement a procedure that reliably identifies negative responders before the stimulation treatment.

While there is a large body of literature on adverse side-effects [41], [44]–[46], NIBS studies typically only consider inter-subject variability in terms of positive- and non-responders, bypassing the potential ethical challenges posed by negative responders. This reservation is arguably a result of the limited understanding of the causes of inter-subject variability in NIBS. Hypothesized explanations for inter-subject variability include non-apparent variations in experimental setups [11], [47], [48], individual differences in brain anatomy [48]– [50], and brain-state dependent susceptibility to NIBS [34], [35], [51], [52]. Factors that have been studied empirically include priming [53], [54], prior activity [55], [56], age [57], [58], attention [59], sex [60], pharmacological influences [61]–[63], genetic variations [64], [65], and the time of day [34]. There is presently no set of factors known, however, that enables a reliable identification of subjects, who do not respond to or may even be harmed by NIBS, before the stimulation treatment [66].

In this article, we demonstrate that inter-subject variability, as measured by the effect of *γ*-range (70 Hz) tACS over contralateral motor cortex on movement speed during a three-dimensional reaching task, encompasses subjects with positive- as well with negative behavioral effects that are sufficiently strong to reach statistical significance on the single-subject level. This result establishes that negative stimulation response is a serious concern for the ethical use of NIBS. We then proceed to show that electrophysiological signatures of resting-state brain activity can be used to predict individual subjects’ stimulation response with high accuracy. Specifically, we present a machine learning pipeline that takes Electroencephalographic (EEG) resting-state recordings of individual subjects as input, and outputs their predicted stimulation response. We apply this pipeline to learn a stimulation response predictor for the present motor performance study, and demonstrate in a prospective study that the stimulation response predictor successfully stratifies subjects into a responder and a non-responder group with statistically significant differences in stimulation effects. In particular, our stimulation response predictor correctly identifies 16 out of 16 subjects who do not respond positively to the stimulation treatment. We then show that successful stimulation response prediction is contingent on resting-state brain signatures recorded directly before NIBS. This finding supports the interpretation that stimulation response is a state and not a trait, i.e., subjects’ susceptibility to brain stimulation may vary over time.

By providing a principled approach to identify subjects, who do not benefit from or may even be harmed by NIBS, before the stimulation treatment, our work constitutes an essential step towards an effective and ethical application of NIBS in clinical settings. Because our results indicate that subjects’ susceptibility to NIBS is a state and not a trait, administering stimulation treatments only when subjects are in a suitable state of mind may further enlarge the range of subjects who benefit from NIBS.

## II. Material and methods

This study conformed to the Declaration of Helsinki, and the experimental procedures involving human subjects described in this paper were approved by the Ethics Committee of the Medical Faculty of the Eberhard Karls University of Tübingen. Informed consent was obtained by all participants, prior to their participation to the study.

### A. Experimental paradigm and data

Each participant attended two sessions, subsequently termed days, separated by a one-day break. On each of the two days, participants were seated on a chair in the middle of four infrared motion tracking cameras (Phase Space, San Leandro, California, USA), facing a visual feedback screen (35′′) at a distance of 1.5 meters, while wearing a specially designed glove with three LEDs on its top for real-time tracking of their arm location. The position of the arm was depicted on the screen in real time as a 3D sphere (cf. Figure 1).

**Fig. 1:**
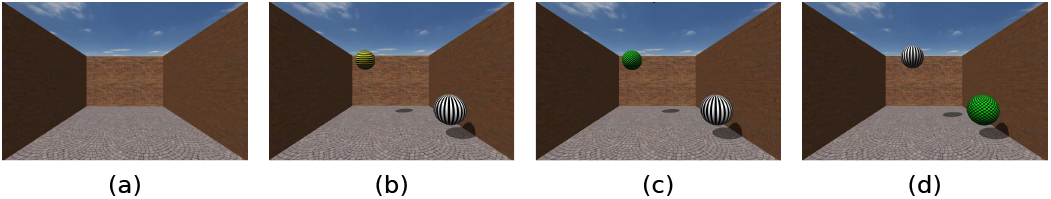
Phases of a trial: a) Subjects wait for the next target. b) A yellow target appears at a random location. Subjects wait for the go-cue, with their current hand position indicated by a white ball. c) A change of target color to green instructs subjects to initiate the reaching movement. d) Subjects have to move their hand back to the starting position, indicated by the green ball.

The experimental paradigm was a 3D-reaching task. In each trial, a target sphere appeared in a simulated 3D space at a random location. The subjects’ goal was to reach the target with their right wrist by moving their arm. Each trial started with a baseline of 5 s, followed by 2.5–4 s during which the target appeared on the screen as a yellow sphere. During this period, subjects had been told to plan but not yet initiate their movement. After the target sphere changed to green, subjects had 10 s to move their arm to reach the target. After a successful reach, a score screen, indicating the movement’s quality, appeared for 2 s. This score was computed as an inverse mapping of their movement’s normalized averaged rectified jerk score (NARJ) to a scale from 0 to 100 [67], [68]. In the last phase of each trial, the sphere appeared at the original starting position of the subjects’ wrist. The trial completed when subjects returned their wrist to the original starting position. If the reach was not completed within 10 s, or if the subject moved before the sphere turned green, the trial was excluded from further analysis.

#### Session 1

During the first day, only electroencephalographic (EEG) (124 active electrodes at 500 Hz sampling rate, BrainAmp DC, Brain Products, Gilching, Germany) and motion tracking (sampling rate of 960 Hz) data were recorded in parallel to the reaching task. The recording session consisted of three blocks of 50 trials each (cf. Figure 2a). Before and after each block, 5 min of resting-state EEG were recorded, during which subjects were asked to relax without moving, focus their eyes on a cross on the screen, and keep their arm in a comfortable position on top of their leg.

**Fig. 2:**
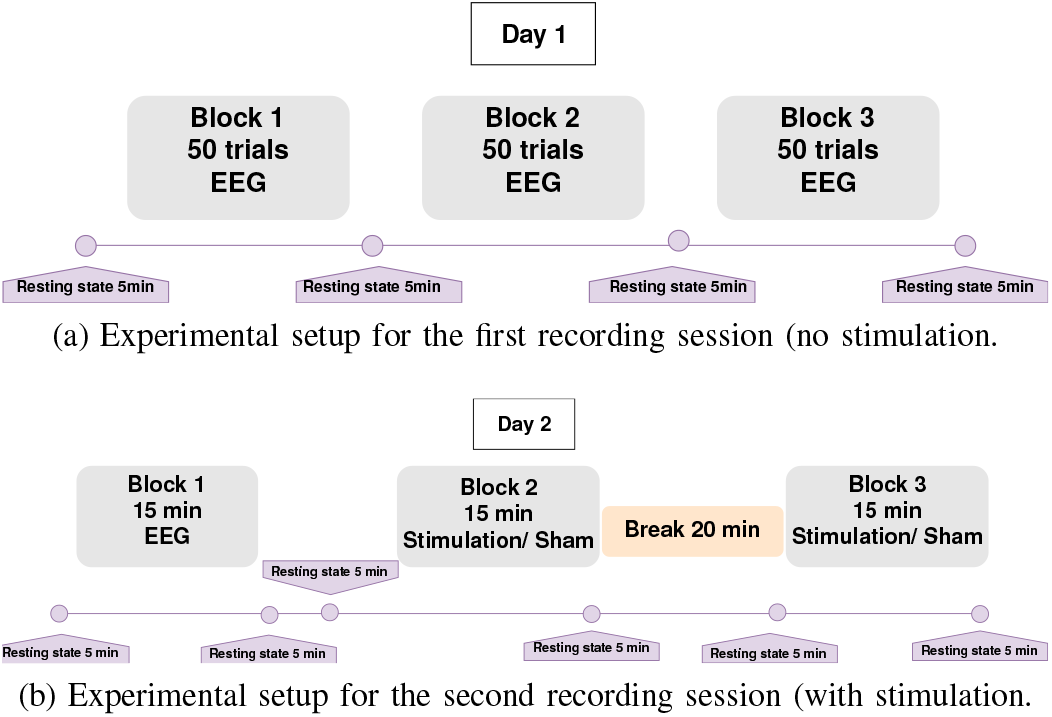
Explanatory diagrams with the EEG and tACS blocks during the recordings of the two experimental sessions, during which each subject participated.

#### Session 2

The second recording day was a crossover randomized stimulation session. This session consisted of three blocks of reaching trials of 15 min each (cf. Figure 2b). During the first block, EEG and motion tracking data were recorded but subjects were not yet stimulated. During the second and third block real- and sham high-definition (HD) transcranial alternating current stimulation (tACS) was applied, respectively, in a randomized order that was not revealed to the subject. A break of 20 min was introduced between the second and the third block, during which the subject was asked to stay seated and relaxed, to avoid carry-over effects. At the end of the session, the participant completed a questionnaire to evaluate the sensation of the stimulation and potential side effects. Before and after each block, a 5 min resting-state EEG was recorded, during which subjects were asked to relax without moving, focus their eyes on a cross on the screen, and keep their arm in a comfortable position on top of their leg.

#### Stimulation setup

We employed a 4 × 1 HD-tACS setup (DC Stimulator Plus, neuroConn GmbH, Ilmenau, Germany) to increase spatial focality relative to the more commonly used two-electrode setup [69]. The equalizer extension box of the DC Stimulator Plus was used to extend the two ordinary square sponge electrodes into a 4 × 1 set of round rubber electrodes of 20 mm diameter (3 cm^2^), with the one electrode in the center, and the four electrodes in a square at a distance of 7.5 cm from the central electrode. The central electrode was placed over channel C3 (left primary motor cortex - M1) and the four surrounding ones over Cz, F3, T7, and P3 [70]. Real stimulation was delivered at 70 Hz and 1 mA peak-to-peak amplitude, while sham stimulation was delivered at 85 Hz and 1 uA peak-to-peak amplitude.

#### Subjects

In the first part of this study (experiments of the first three months) twenty healthy, right-handed subjects participated. One subject did not attend the second day of recordings and was excluded from further analyses. The remaining 19 subjects (nine female) had an average age of 28.37 years with a standard deviation of 8.57 years. In the second part of the study (experiments of the last three months) twenty-two new subjects were recorded and stimulated following the exact same protocol. This second group was only used as an evaluation group, as described in detail in subsection II-E].

### B. Analysis of behavioral data

We quantified the behavioral effect size of *γ*-tACS over contralateral M1 by the difference between the average reaching speed during the real- and the sham-stimulation block, because *γ*-tACS over motor cortex has been reported to influence movement velocity [21], [34], [71]. Furthermore, arm velocity has been considered one of the most stable variables related to the motor cortex activity [72], [73]. To compute the effect sizes on the level of individual subjects, we first computed, for every trial and subject, the trial-averaged reaching speed. This was done by, first, identifying the part of each trial which corresponded to the subject’s movement, i.e., from the “Go” phase until the reaching of the target. We then extracted the *x*, *y* and *z* coordinates from the frames of the camera and calculated the mean velocity as the amplitude of the discrete positional derivative [68]. For each subject, we computed the block-averaged reaching velocities by averaging the trial-averaged velocities within the real- and the sham-stimulation block. If a trial-averaged velocity deviated from the block-averaged velocity by more than three standard deviations the trial was rejected as an outlier. Finally, we computed the difference between the block-averaged velocities of the real- and the sham-stimulation blocks and normalized the difference by the standard deviation of each subject’s sham-stimulation block to obtain the subject-level behavioral effect sizes. To compute the group-level behavioral effect size, we averaged the subject-level effect sizes and normalized by their standard deviation.

To test for a statistically significant behavioral stimulation effect on the level of individual subjects, we performed, for each subject, a two-sided, t-test on the trial-averaged arm velocities of the real- and the sham-stimulation block. We built the null-distribution by randomly permuting the assignment of trials to the real- and sham-stimulation block 10.000 times. After every permutation, we re-computed the subject’s average speed difference between the real- and the sham-block. We calculated the *p*-value as the frequency at which samples from the null-distribution exceeded the original absolute average speed difference between the real- and the sham-block. Subjects with *p <* 0.05 and larger average speed during the real-compared to sham-block were subsequently termed *responders*. The remaining subjects, who consist of those with non significant *p*-values, and those with significant reduction of arm speed (to whom we may also refer as “negative responders”) were termed combined *non-responders*.

To test for a statistically significant behavioral stimulation effect on the group-level, we performed a two-sided, paired t-test on the single-subject effect sizes. We built the null-distribution by randomly flipping every subject’s block-average velocities between the real- and sham-blocks 10.000 times. After random permutation, we re-computed the group-level behavioral effect size as described above. We calculated the *p*-value as the frequency at which samples from the null-distribution exceeded the original absolute group-level effect size.

### C. Analysis of EEG data

We first cleaned each subject’s EEG data from non-cortical artifacts by Independent Component Analysis (ICA), and then computed resting-state bandpower in canonical EEG frequency bands.

For each subject and session, we concatenated the raw data of all resting-state recordings, high-pass filtered the data with a Butterworth filter at 3 Hz, and re-referenced the data to common-average reference. We then used the SOBI algorithm [74] to extract 64 independent components (ICs). We manually inspected the topography of every IC, and discarded those ICs that did not show a cortical topography [75]. The remaining cortical ICs (ranging between four and 18 across subjects) were re-projected to the scalp level, and the individual resting-state recordings were reconstructed. For each subject, resting-state, and electrode, we normalized the data by z-scoring. The reason for the few kept ICs was that during the manual artefact cleaning we were particularly cautious not to include contaminated signals. Therefore, we were conservative by keeping only IC components that looked cortical, rejecting ambiguous ones.

For every subject, resting-state, and electrode, we then computed log-bandpower in eight canonical EEG frequency bands: high *δ* (3–4 Hz) (due to 3Hz high-pass filter), *θ* (4–8 Hz), *α* (8–12 Hz), *β* (12–25 Hz), *γ*_1_ (25 –45 Hz), *γ*_2_ (45–65 Hz) (although line power at 50 Hz is included), *γ*_3_ (65–90 Hz), and *γ*_4_ (90–160 Hz). This was done by windowing the data with a Hamming window, computing the Discrete Fourier Transform, taking the average of the absolute values of all frequency components within each of the eight frequency bands, and finally taking the natural logarithm.

### D. Training of the stimulation response predictor

We trained a linear discriminant analysis (LDA) classifier to predict subjects’ category (*responder* vs. *non-responders*) from their resting-state EEG. Due to the small sample size, we selected two EEG channels over left-(CCP3h) and right motor cortex (CCP4h) and one channel over parietal cortex (Pz) as input to the classifier (channel C3 directly over left motor cortex was blocked by the stimulation electrodes, hence for symmetry we did not use C4 as well). For each of the eight frequency bands (cf. Section II-C) and the three resting-state recordings on the stimulation day, we evaluated the prediction accuracy of the classifier by leave-one-subject-out cross-validation. We tested the statistical significance of each of the 24 settings (eight frequency bands times three resting-states) by a permutation test with 1000 permutations. To build the null-distribution, we randomly permuted the labels on the training set of each cross-validation fold, retrained the classifier, and classified the subject in the test set. We calculated the *p*-value as the frequency at which samples from the null-distribution exceeded the original prediction accuracy. We then selected the best-performing combination of frequency-band and resting-state to train the final stimulation response predictor (SRP) on all 19 subjects.

### E. Validation of the stimulation response predictor

To validate the SRP, we recruited 22 new subjects (eleven female, average age of 26.81 years with a standard deviation of 6.32 years). We employed the same EEG processing pipeline as for the first group of subjects (cf. Section II-C), except that we used the EEG data of the first session (recorded on day one) to compute the ICA. To clean the EEG of the stimulation session (recorded on day two) from non-cortical artifacts, we applied the spatial filters derived on day one to the EEG data of day two, and reprojected only those ICs that corresponded to cortical sources on day one. In this way, we minimized the probability that any manual selection of ICs on day two could have confounded the predictions of the SRP. We then applied the trained SRP, as described in Section II-D, to the first resting-state recording of every subject in the validation group, and compared the predicted categories with those derived from the behavioral analysis described in Section II-B (the categorization of subjects in the validation group into *responders* and *non-responders* is shown in Table II in the Appendix).

To test for a statistically significant difference in the behavioral stimulation effect between the predicted responder and non-responder group, we employed a one-sided permutation-based t-test: We randomly permuted the predicted assignments of subjects to the responders- and non-responders group 10.000 times. After every permutation, we recomputed the group-level effect size within each group (responders vs. non-responders) as described in Section II-B, and calculated the *p*-value as the frequency at which the permuted difference in effect sizes exceeded the original one.

### F. Association of stimulation response with external factors

We tested the stimulation response of individual subjects (*responders* vs. *non-responders*, cf. Section II-B) for associations with four external factors: gender, order of the sham- and real-stimulation block, block of the strongest perceived sensation of stimulation as reported by the subjects, and baseline motor performance during the first session. For all analyses, we pooled all 41 subjects from the training- and the validation group.

To test for an association of the stimulation response with gender (female, male) and order of the stimulation block, respectively, we used Fisher’s exact test. To test for an association of the stimulation response with the block of the strongest sensory sensation, we employed a chi-squared test. Finally, to test for an association between the average reaching speed during the first session (on day one) with subsequent stimulation response in the second session (on day two), we performed an ANOVA for the average movement speed across all three blocks on day one. For all tests, we chose a significance level of *α* = 0.05.

### G. Signal to noise ratio of EEG in high gamma range

To examine and eliminate the possibility that our findings in the high gamma range are coincidental or mostly noise, we performed the following analysis. To make sure that we do not introduce any bias during the manual artefact correction process, we tested the statistical association between the number of kept ICs and the response group. Moreover, we examined the association between the type of the kept ICs (i.e. exhibiting a horizontal or vertical dipole) and the response group (all the kept ICs can be found in the supplementary material). Finally, we estimated the frequency-dependent signal-to-noise (SNR) ratio of our EEG amplifier by comparing the EEG resting-state recordings with the amplifier’s noise floor, measured by submerging the EEG electrodes in a saline solution. Results are presented in Figure 9 and discussed in subsection IV-C.

## III. Results

### A. Positive- and negative stimulation effects in tACS

We assessed inter-subject variability of the behavioral response to tACS in a motor performance study with a cross-over design. Nineteen healthy subjects performed 15-minute blocks of 3D reaching movements with their right hand to targets appearing at random locations in a 2D visual feedback setup (Figure 1).

Figure 3 shows the histogram and estimated probability density function (Gaussian kernel density estimate with a kernel width of 0.5 [76], [77]) of effect sizes across subjects. While the group-level effect size of 0.33 is not statistically significant (*p* = 0.1018, subject-level permutation-based paired t-test, cf. Section II-B), we found a substantial variation in the effect sizes of individual subjects, ranging from −1.12 to 1.51 with a standard deviation of 0.72. Statistical tests for significant effect sizes at the individual subject level revealed seven subjects with a statistically significant *positive* and four subjects with a statistically significant *negative* effect size (at significance level *α* = 0.05, two-sided trial-level permutation-based t-test). The remaining eight subjects did not show a statistically significant effect at the individual subject level (individual *p*-values are shown in Table I in the Appendix). These results demonstrate that *γ*-tACS can have positive-as well as negative behavioral effects on motor performance, which poses an ethical challenge to tACS studies ^1^. In the following section, we demonstrate how to address this challenge by predicting individual subjects’ stimulation response from their resting-state configuration of brain rhythms.

**Fig. 3:**
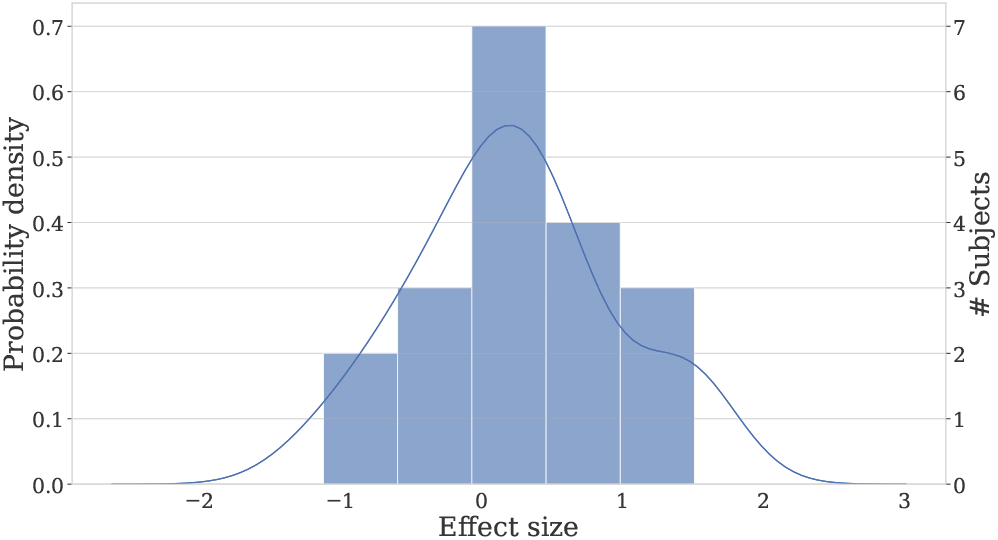
Histogram and estimated probability density function of stimulation response.

**TABLE I:** Categorization of subjects into *responders* and *non-responders*, first group of recordings. ΔVelocity *>* 0 refers to subjects with a higher movement speed in the real- vs. the sham-stimulation block.

| Subject | *p*-value | $\Delta\text{Velocity} > 0$ | Category |
| --- | --- | --- | --- |
| 1 | 0.0005 | 1 | responder |
| 2 | 0.6217 | 1 | non-responder |
| 3 | 0.0902 | 1 | non-responder |
| 4 | 0.0121 | 0 | non-responder |
| 5 | <0.0001 | 1 | responder |
| 6 | 0.9318 | 1 | non-responder |
| 7 | 0.3979 | 1 | non-responder |
| 8 | 0.1628 | 1 | non-responder |
| 9 | 0.9220 | 0 | non-responder |
| 11 | 0.0154 | 0 | non-responder |
| 12 | 0.0251 | 1 | responder |
| 13 | 0.0384 | 0 | non-responder |
| 14 | 0.0800 | 0 | non-responder |
| 15 | 0.8484 | 1 | non-responder |
| 16 | <0.0001 | 1 | responder |
| 17 | <0.0001 | 0 | non-responder |
| 18 | 0.0077 | 1 | responder |
| 19 | 0.0370 | 1 | responder |
| 20 | 0.0455 | 1 | responder |

**TABLE II:** Categorization of subjects into *responders* and *non-responders*, second (validation) group of recordings. ΔVelocity *>* 0 refers to subjects with a higher movement speed in the real- vs. the sham-stimulation block.

| Subject | *p*-value | $\Delta\text{Velocity} > 0$ | Category |
| --- | --- | --- | --- |
| 21 | 0.0001 | 0 | non-responder |
| 22 | 0.0004 | 1 | responder |
| 23 | 0.0119 | 1 | responder |
| 24 | 0.0093 | 1 | responder |
| 25 | 0.0353 | 1 | responder |
| 26 | 0.5704 | 1 | non-responder |
| 27 | 0.0001 | 1 | responder |
| 28 | <0.0001 | 0 | non-responder |
| 29 | 0.5334 | 0 | non-responder |
| 30 | 0.5449 | 0 | non-responder |
| 31 | 0.6537 | 1 | non-responder |
| 32 | 0.8046 | 0 | non-responder |
| 33 | 0.8770 | 1 | non-responder |
| 34 | 0.0055 | 0 | non-responder |
| 35 | 0.0678 | 0 | non-responder |
| 36 | 0.6119 | 1 | non-responder |
| 37 | 0.2996 | 0 | non-responder |
| 38 | 0.0048 | 1 | responder |
| 39 | 0.4444 | 1 | non-responder |
| 40 | 0.7515 | 1 | non-responder |
| 41 | 0.5464 | 0 | non-responder |
| 42 | 0.0075 | 0 | non-responder |

### B. Resting-state EEG predicts tACS stimulation response

Before and after each block, we recorded a high-density resting-state electroencephalogram (EEG), manually cleaned the data from non-cortical artifacts, and computed subject-specific log-bandpower estimates for all channels in canonical EEG frequency bands ranging from 1–160 Hz (the EEG analysis is described in detail in Section II-C). We then separated subjects into those with a statistically significant positive stimulation response, subsequently called the *responders*, and the remaining subjects (both negative responders and subjects with no significant response), subsequently termed the *non-responders* (cf. Section II-B). The reason why we did not use three different groups in our prediction pipeline (“positive”, “negative” and “non-responders”) was the limited number of subjects for training such a classifier. ^2^ Therefore, we proceed our analysis with the aforementioned two groups.

The group-average topography of power in the *γ*-range (90–160 Hz), recorded prior to the first (real- or sham-) stimulation block, revealed strong resting-state *γ*-power over contralateral motor cortex in the *responders* but not in the *non-responders* (Figure 4). This result suggests that only those subjects, who already had strong *γ*-power over contralateral motor cortex before the start of the stimulation, showed a subsequent positive behavioral response to contralateral *γ*-tACS. To systematically evaluate the predictive value of resting-state brain rhythms for stimulation response, we trained a machine learning algorithm to predict individual subjects’ responses to *γ*-tACS from their resting-state brain rhythms. Specifically, we selected three EEG channels over left motor, right motor and central parietal cortex, computed the log-bandpower during the resting-state recorded prior to the first stimulation block in canonical frequency bands, and then employed a leave-one-subject-out cross-validation procedure to assess the ability of a linear discriminant classifier (LDA) to predict each subject’s group (responder vs. non-responder).

**Fig. 4:**
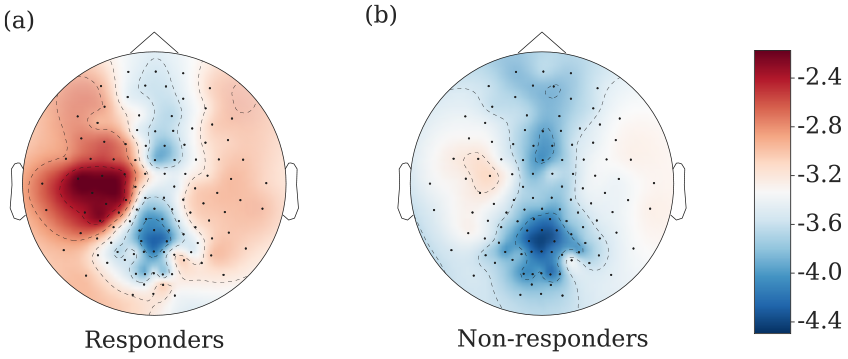
Group-averages for responders (a) and non-responders (b) of high *γ* (90–160 Hz) log-bandpower during the resting state recorded at the end of the first block of the second session (prior to stimulation blocks). Each subject’s data were z-scored before the across-subjects averaging.

The details of the prediction pipeline are described in Section II-D. We found the prediction accuracy to increase with frequency, peaking at 89.47% in the band from 90–160 Hz (*p <* 0.001, permutation test, cf. Section II-D for details). Prediction accuracies and *p*-values for all frequency bands are shown in Figure 6 in the Appendix. To test the robustness of the prediction pipeline, we repeated the machine learning procedure for all three resting-state recordings preceding the first stimulation block. We found that all resting state recordings enabled above chance level prediction in the 90–160 Hz band. Prediction accuracies of the remaining bands varied across the different resting states.

These results are an indication that high-gamma power can carry important information for the prediction of subjects’ behavioral response to *γ*-tACS over contralateral motor cortex.

### C. Subject stratification by resting-state EEG enhances effect sizes

To evaluate the practical utility of the response stratification pipeline described in the previous section, we performed an additional validation study with 22 new participants. Based on the results described in the previous section, we chose the classifier trained on the resting-state recorded after the first block in the 90–160 Hz frequency band for the validation study. This classifier was trained on the first group of subjects and then used out-of-the-box to predict the stimulation response for each subject in the validation group from a resting-state EEG recorded prior to the first block of trials (see Section II-E for details).

In the validation group, we observed a group-level behavioral effect size of 0.12 (*p* = 0.2847, subject-level permutation paired t-test), with subject-level effect sizes ranging from −0.94 to 1.19 (see Figure 7 in the Appendix). The true response groups of the validation subjects based on their performance can be found in Table II in the Appendix. We used the pre-trained SRP to classify the subjects on the two response groups (responders and non-responders). The EEG-based stratification of subjects resulted in group-level effect sizes of 2.46 and −0.17 for the predicted responders and non-responders, respectively, a statistically (*p* = 0.0048, one-sided permutation-based t-test) and practically highly significant difference. In particular, all four subjects with a statistically significant negative- and all 12 subjects with no statistically significant stimulation response were correctly classified by SRP to the group of non-responders. Further, only two subjects with a statistically significant positive stimulation response were mis-classified as non-responders. These behavioral results are summarized in Figure 5. The group-averaged topographies of log-bandpower in the *γ*-range of the predicted responders and non-responders, which closely resemble those observed in the training group shown in Figure 4, are displayed in Figure 8 in the Appendix.

**Fig. 5:**
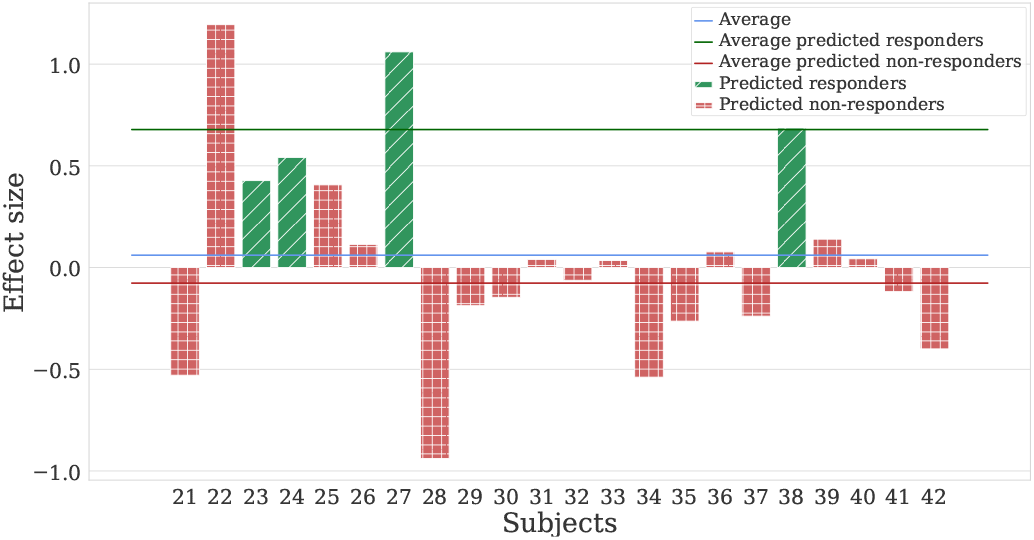
Individual effect-sizes and predicted stimulation responses in the validation group. The height of the bars indicate the effect size of the real stimulation for each subject. The two colours indicate the two classes (“Reponderes” and “Non-responders” as have been defined in Section II-B) that our SRP predictor classifies the subjects from their resting state EEG. As we can see, comparing our predictions with the true behavioural response groups in Table II, our predictor correctly identifies all the “Non-responders” and four “Responders”. It only misses two true positive responders (subjects 22 and 25).

**Fig. 6:**
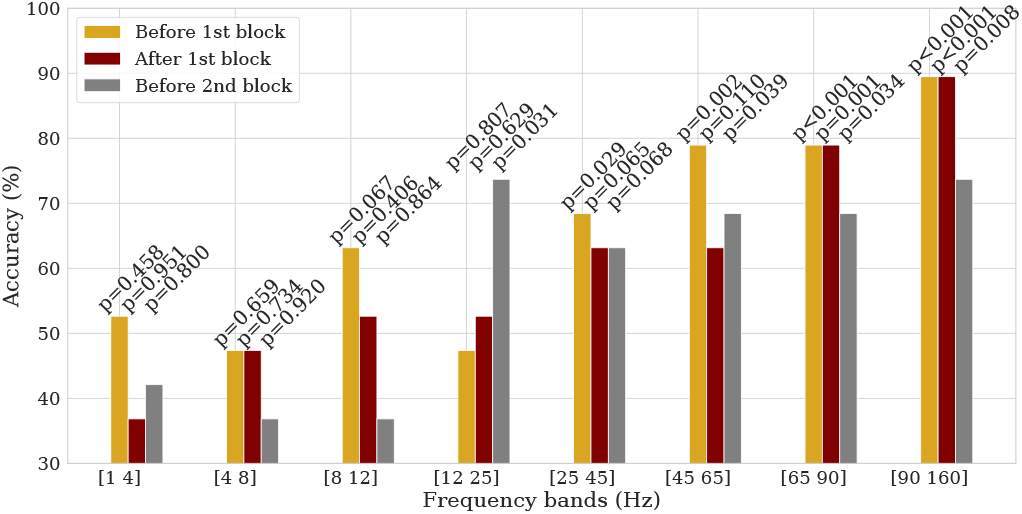
Leave-one-subject-out cross-validated prediction accuracy of stimulation response in the first group of subjects across canonical frequency bands and resting-states, cf. Section II-D for details. Stimulation day (day 2).

**Fig. 7:**
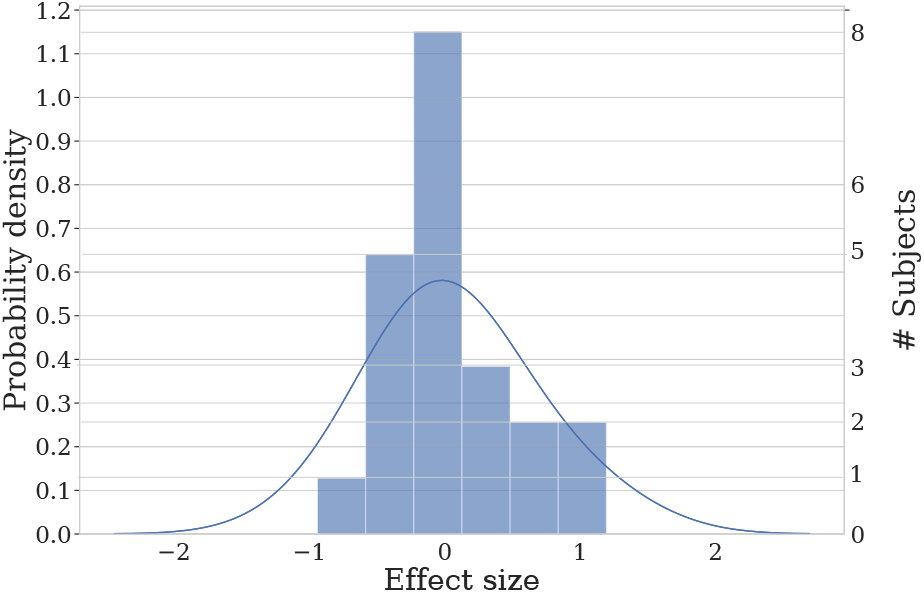
Histogram and estimated probability density function of stimulation response in the validation group.

### D. Stimulation response is contingent on brain state

In a next step, we employed the validated prediction pipeline to test whether stimulation response is a state or a trait, i.e., whether subjects’ response to *γ*-tACS changes or remains invariant over time. To do so, we pooled all 41 subjects, trained our prediction pipeline on the *γ*-power (90–160 Hz) of each of the four EEG resting states of their first session, i.e., two days before the stimulation session, and predicted the stimulation response in the second session with leave-one-subject-out cross-validation (all other settings were identical to those described in Sections II-C and II-D). A statistically significant prediction accuracy in this setting would imply that the configuration of subjects’ brain rhythms is also predictive for their stimulation response two days later. We did not, however, find any evidence in favor of this conclusion. Instead, training on brain activity of the first recording session resulted in statistically non-significant prediction accuracies between 62.5% and 68.3% (p-values of 0.23, 0.24, 0.27 and 0.73 for the four resting states of day one). This observation implies that stimulation response is contingent on subjects’ brain-state directly prior to the stimulation, i.e., subjects’ stimulation response is a state and not a trait.

Applying the original stimulus-response predictor, as described in Section II-E, to the resting-state recordings of the first day, we estimate that out of the 28 subjects, who did not respond positively to the stimulation on day two, five subjects would have responded positively on day one (prediction results for individual subjects are shown in Table III in the Appendix). As such, the percentage of subjects, who can benefit from tACS, may increase if they are stimulated at the right time.

**TABLE III:** Subjects’ predicted behavioral response from resting-state EEG data recorded on day one versus actual behavioral response measured on day two.

|  |  |
| --- | --- |
| Subjects predicted as responders on day one | 15, 18, 19, 22, 23, 24, 27, 33, 34, 36, 38, 42 |
| Actual non-responders on day two | 2, 3, 4, 6, 7, 8, 9, 11, 13, 14, 15, 17, 21, 26, 28, 29, 30, 32, 33, 34, 35, 36, 37, 39, 40, 41, 42 |
| Intersection | 15, 33, 34, 36, 42 |

## IV. Discussion

### A. Contributions

Our results demonstrate that resting-state signatures of human brain rhythms, recorded prior to NIBS, can distinguish responders from non-responders with high accuracy. This contribution is essential for a safe and ethical application of NIBS in research and treatment. As it has been reported that NIBS can have behavioral effects of opposite polarity relative to the intended stimulation effect in individual subjects (cf. Section III-A), a reliable exclusion criterion for subjects with either a negative stimulation response or no response at all, ensures that no subjects are harmed and that redundant stimulation sessions are avoided. This issue is of particular relevance in clinical settings, where NIBS is employed to cause long-term and, possibly, permanent changes, and where the stimulation sessions are expensive for patients with no potential of benefiting from it.

### B. The task

Here we briefly discuss our rationale for the selection of the specific task. The reason why we chose this task is because it allows for a free 3D arm movement that mimics natural reaching movements, which we would like eventually to enhance and facilitate through stimulation. We are interested in the arm speed as our behavioural metric, because this metric has been found to be the most robust variable related to the motor cortex [73], with Moran and Schwartz [72] having even proposed a canonical model for it. In addition to the aforementioned reason, we preferred as behavioural metric the speed over the normalized average rectified jerk, which we report as score to the subjects, because NARJ is the second derivative of the speed, which already accumulates a lot of error in the measurement, starting from the position recorded from the infrared cameras. Therefore, we selected this task as it could help us measure the arm speed in a 3d movement task that seems natural. For the present study, we are not interested in the performance of the subjects in terms of successfully reached targets as a metric, as this would potentially include a more complex cognitive procedure. This is the reason why we focus in the relation between the motor cortex activity and the arm speed.

### C. High frequency band carries information about the response to gamma motor stimulation

Being aware of the fact that analysis of high-gamma EEG power are controversial, at first, we were surprised to find that such a high frequency (90–160 Hz) power was the most predictive of tACS response. Here we discuss why, based on our analysis, we can be optimistic that this is not an artefact or noise-driven finding. First, all predictions were made from resting-state EEG recordings without tACS stimulation. As such, the high-gamma power features, that we used for predicting the behavioral responses, could not stem from stimulation artifacts. Moreover, we employed a particularly conservative IC rejection procedure, following the recommendations by [75], to minimize the probability of retaining artifactual sources in the measurements. This approach also explains why we kept a few components per subject. To make sure that we did not introduce any bias during this manual process, we also tested the statistical association between the number of kept ICs and the response group of each subject. Kruskal Wallis test yielded no significant association (p = 0.23). Furthermore, no association was found between the type of the kept ICs (i.e. exhibiting a horizontal or vertical dipole) and the response group.

In our preliminary analysis, we have tested frequencies up to 200 Hz to assure that they are not predictive. No frequencies above 160*Hz* were found to be predictive. Moreover, the grand-average topoplots (Figures 4 and 8) indicate that high gamma power in our study was strongest over contralateral motor cortex, which indicates that contamination by EMG activity, which would be strongest over frontal- and occipital areas [75], was also not an issue here.

**Fig. 8:**
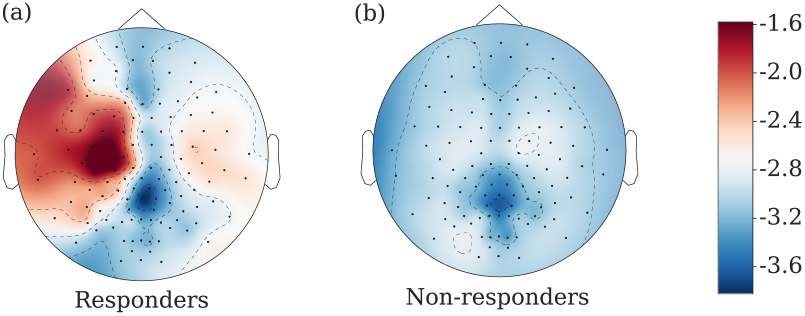
Group-averages for predicted responders (a) and non-responders (b) in the validation group of high *γ* (90–160 Hz) log-bandpower during the resting state recorded at the beginning of the stimulation day (prior to stimulation blocks).

Finally, to provide further evidence that EEG measurements in the high gamma range are feasible, we estimated the frequency-dependent signal-to-noise (SNR) ratio of our EEG amplifier by comparing the EEG resting-state recordings of the three channels used by the classifier with the amplifier’s noise floor, measured by submerging the EEG electrodes in a saline solution, as described in subsection II-G. Figure 9 in the Appendix indicates an average SNR between 6 and 7 dB in the 90 to 160 Hz range, i.e., the recorded signal in the high gamma range is at least four times stronger than the inherent measurement noise. This observation is in line with recent work that indicates that the feasibility to measure human gamma power in scalp EEG is not limited by recording hardware but rather depends on subjects’ morphology [78].

**Fig. 9:**
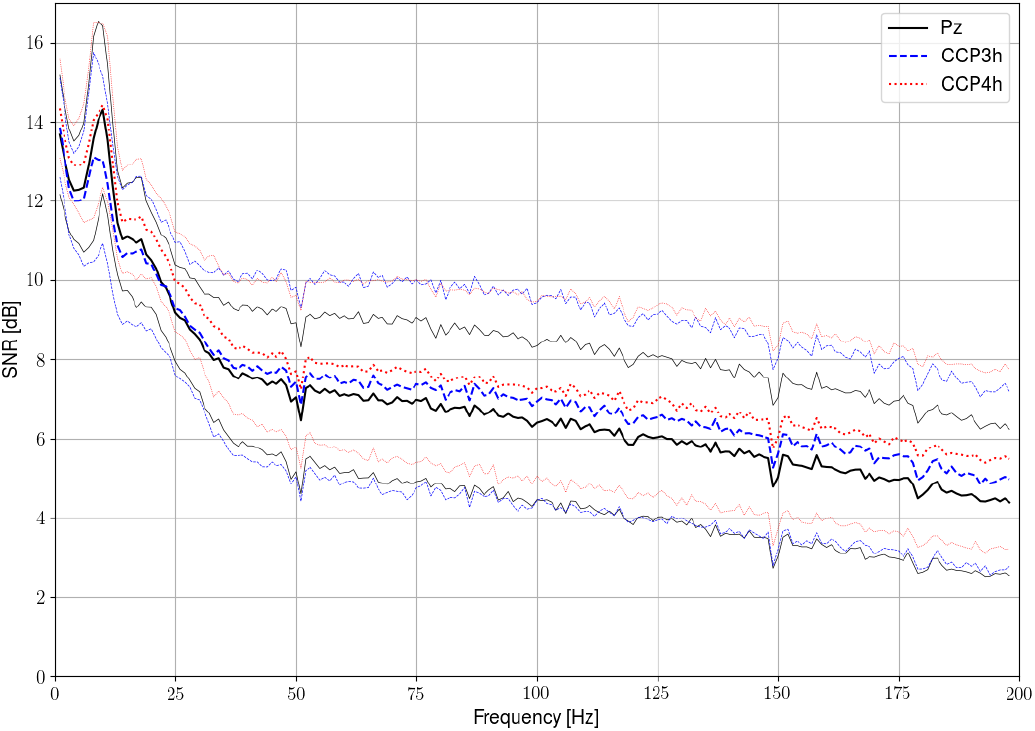
Mean SNR plus, minus one standard deviation across subjects’ raw resting-state data (i.e., no common average and no ICA cleaning) during the 5 min of resting state used by our SRP classifier (beginning of 2nd block, Session 2), with respect to the noise floor measured for the same time length, by submerging the electrodes in a saline solution. The three different colours/textures refer to the three channels used by the SRP classifier (CCP3h, CCP4h and Pz). The bold lines refer to the mean SNR accross subjects, while the faded lines depict one standard deviation. As we can seethe actual EEG has more power than the noise floor in the high gamma range, which is another indication that our classifier is not based on noise.

Therefore, it seems plausible that high gamma range indeed carries such significant information.

### D. Plausible neurophysiological mechanism

In our experiments we found strong resting-state *γ*-power over contralateral motor cortex to be indicative of a positive stimulation response to tACS (in terms of arm speed). As we outline in the following, this finding is in line with our current understanding of the neurophysiological effects of *γ*-tACS and the role of *γ*-power in fronto-parietal networks for motor performance [79]. Resting-state *γ*-power in primary motor cortex positively correlates with *γ*-aminobutyric acid (GABA) levels [80]–[84]. Because *γ*-tACS over motor cortex decreases GABA [85], and decreases in motor cortex GABA levels correlate with increased motor performance [86], high resting-state *γ*-power may signal a brain state in which motor performance can be improved through tACS-induced reduction of GABA levels. Low resting-state *γ*-power, in contrast, would signal a brain state in which GABA levels are already low, thus limiting the extent of potential further reduction by *γ*-tACS. We note that this explanation is also in line with our finding that stimulation response is contingent on the current brain state (cf. Section III-D). While our argument is consistent with the neurophysiological GABA activity, it is worth mentioning that there are studies on macaques, that exhibit enhancing of the ongoing gamma activity by gamma tACS [7], [8]. Nevertheless these studies protocols differ significantly from ours in the positioning of electrodes. Krause et al. [7] do not target the motor cortex, but the hippocampus and visual cortex of anaesthetized animals; which could explain the discrepancy with our findings. Finally, Johnshon et al. [8] place the bipolar electrodes over left and right temples, which could also induce different effects from our setup.

### E. Evaluation of external factors’ role in behavioural response to stimulation

To further probe the state vs. trait hypothesis, and to examine the possibility that behavioural response might be affected by possible differences in sensation between sham and real stimulation, we tested a range of subject traits and experimental factors, including gender, order of real-/sham-stimulation, block of strongest-reported stimulation sensation and behavioral performance on the first day of the experiment, for associations with stimulation response. None of these factors reached a statistically significant association (Fisher’s exact test, Chi-squared test and ANOVA, see Section II-F). This observation is in line with previous work on explaining individual subjects’ stimulation response (cf. Section I), and underlines the significance of subjects’ current brain state for their stimulation response. Of course, besides the gamma activity there can be other factors that play a hidden confounding role, or that we have not yet observed. Such an analysis would require a causal inference approach that is robust against hidden common causes, such as [87]–[91], or even a causal-effect attribution approach [92] which is beyond the scope of this paper. Nevertheless, we are confident in excluding the possibility that the measured behavioural response is caused by different sensation between the two conditions for the following reason. Subjects evaluated a range of possible sensations (i.e. tingling, burning, phosphates, itching etc.) for both conditions, without knowing which condition was what, and the statistical analysis showed no significant association of the response group, neither with the block of strongest reported sensation nor with any of the reported sensations.

### F. Outlook

Our results further indicate that the percentage of subjects, who can benefit from NIBS, may be increased when subjects are stimulated at the right time. Concurrently, the neurophysiological interpretation of our results raises the question whether the effects of stimulation lie within the range of normal variations in behavioral performance, i.e., whether NIBS induces a beneficial state of mind that can also occur spontaneously, or whether NIBS can enhance behavioral performance beyond subjects’ natural limits. Either way, a natural extension of our stimulation response prediction pipeline would be to consider multiple stimulation settings that vary in parameters such as spatial and spectral focus, paving the way for personalized NIBS. Being motivated by the exhibited difference in gamma power between the two groups – as depicted in Figures 4 and 8– another future extension would be to combine this pipeline with a pre-stimulation step of gamma-modulation, to study whether a self-modulated rise of the gamma activity could ensure positive response to the tACS.

## V. Conclusion

We believe that identification of responders and non-responders prior to the application of stimulation treatment is an important first step towards personalized brain stimulation. In our work, we show that resting-state high-gamma power prior to stimulation enables this differentiation. Specifically, we demonstrate in a first experimental group of 19 participants that subjects’ resting-state EEG predicts their motor response (arm speed) to gamma (70 Hz) transcranial alternating current stimulation over the contralateral motor cortex. We then ascertain in a prospective stimulation study with twenty-two new subjects that our prediction pipeline achieves a reliable stratification of subjects into a responder and a non-responder group.

## Footnotes

1 Even after strict multiple test correction on the *p*-values there are both positive and negative responders. After performing a Holm-Sidak multiple-test correction at a significance level 0.05, on the 19 p-values reported in Table I, there are still subjects with significant positive and significant negative response. The p-values of these first group after the multiple-test correction can be found below the Table I

2 In a preliminary analysis we did not manage to reach satisfactory classification accuracy for three groups, for none of the frequency bands.

## VII. Appendix

Here we present additional figures from our analysis during the two periods of recordings/stimulations, which further support our findings.

After performing a Holm-Sidak multiple-test correction at a significance level 0.05, the corrected p-values of the first group are: 0.0079 0.9795, 0.4867, 0.1567, 0.0018, 0.9965, 0.9208, 0.6556, 0.9965, 0.1827, 0.2629, 0.3394, 0.4867, 0.9965, 0.0018, 0.0018, 0.1094, 0.3394, 0.3423. As we can see there are still subjects with significant positive (subjects 1, 5, 16) and significant negative response (subject 17).

The corrected p-values are: 0.0021, 0.0075, 0.1543, 0.1307, 0.3732, 0.998, 0.0021, 0.0021, 0.9989, 0.9989, 0.9989, 0.9989, 0.9989, 0.0894, 0.5693, 0.9989, 0.9801, 0.0829, 0.9971, 0.9989, 0.9989, 0.1134, which means that even after the strict multiple testing correction there are still both significantly positive (subjects 2, 7) and significantly negative responders (subjects 1, 6).

